# Cellquant: a vibecoder’s guide to image analysis

**DOI:** 10.64898/2026.03.09.710634

**Authors:** Abani Neferkara, Leah Chaney Winner, Asif Ali, David Pincus

## Abstract

Quantitative fluorescence microscopy is central to modern cell biology, yet experimentalists lacking computational expertise have limited ability to extract reproducible measurements from the images they collect. Here we present `cellquant`, a pipeline to analyze multi-channel fluorescence images that performs cell segmentation, subcellular localization and colocalization analyses, morphometric quantification, and spatial proximity calculations in two or three dimensions. By replacing the requirement for computational expertise with iterative assistance from large language models (i.e., “vibecoding”), `cellquant` enables experimentalists with any level of coding experience to quantify imaging data. We validate `cellquant` on stress granule formation in human HCT116 cells and nucleolar reorganization across a temperature gradient in budding yeast. Dimensionality reduction of a 45-feature output from 8,309 yeast cells resolved cell state transitions and revealed subpopulations of non-responders. Because the `cellquant` interface is text based, the exact command used in any analysis can be recorded and re-executed, generating reproducible quantification with visual quality control and statistical rigor. Code, documentation, and datasets are freely available.

## Introduction

Fluorescence microscopy provides the visual evidence underlying much of modern cell biology. Advances in segmentation algorithms, particularly deep learning approaches such as Cellpose [Stringer et al., 2021, Pachitariu and Stringer, 2022], have largely solved the technical challenge of identifying individual cells in microscopy images. Likewise, there are established algorithms for detecting sub-cellular structures, quantifying colocalization, and calculating distances between cellular compartments.

Despite the availability of these algorithms, experimental cell biologists do not routinely integrate quantitative image analysis into their protocols. Unlike next-generation sequencing, the analysis of which has become semi-standardized, image processing and quantification remains bespoke. Lack of training in basic computational skills like the ability to navigate directories from the command line limits what many biologists will attempt in their analyses. Even for those with fluency, it is often the case that individual tools must be assembled into nontrivial pipelines, requiring expertise not only in image segmentation but also in statistical analysis. Existing tools have substantially reduced the analysis barrier, including CellProfiler [Stirling et al., 2021] via a graphical pipeline builder, Fiji/ImageJ [Schindelin et al., 2012] via macro-based automation, and napari [Sofroniew et al., 2022] via interactive visualization. However, each requires the user to learn how to use new software, and re-configuring these tools for each new biological question still often demands computational expertise. Moreover, although these tools now have artificial intelligence (AI) features, they were not explicitly designed to work alongside coding assistants and large language models (LLMs).

LLM-based assistants enable a new paradigm in which a biologist can describe their analysis in natural language and have a chatbot write, modify, and run code. This approach, colloquially termed “vibecoding,” works well for image analysis, where the ability of an experimentalist to recognize proper segmentation and identify subcellular structures is essential to validate automated pipelines (Figure 1A). To leverage LLM capabilities, existing image processing software packages have integrated AI assistants that can generate bespoke code on demand. While powerful, these tools still require users to already be fluent in specific software and the code itself is rarely standardized, versioned, or vetted.

**Figure 1:**
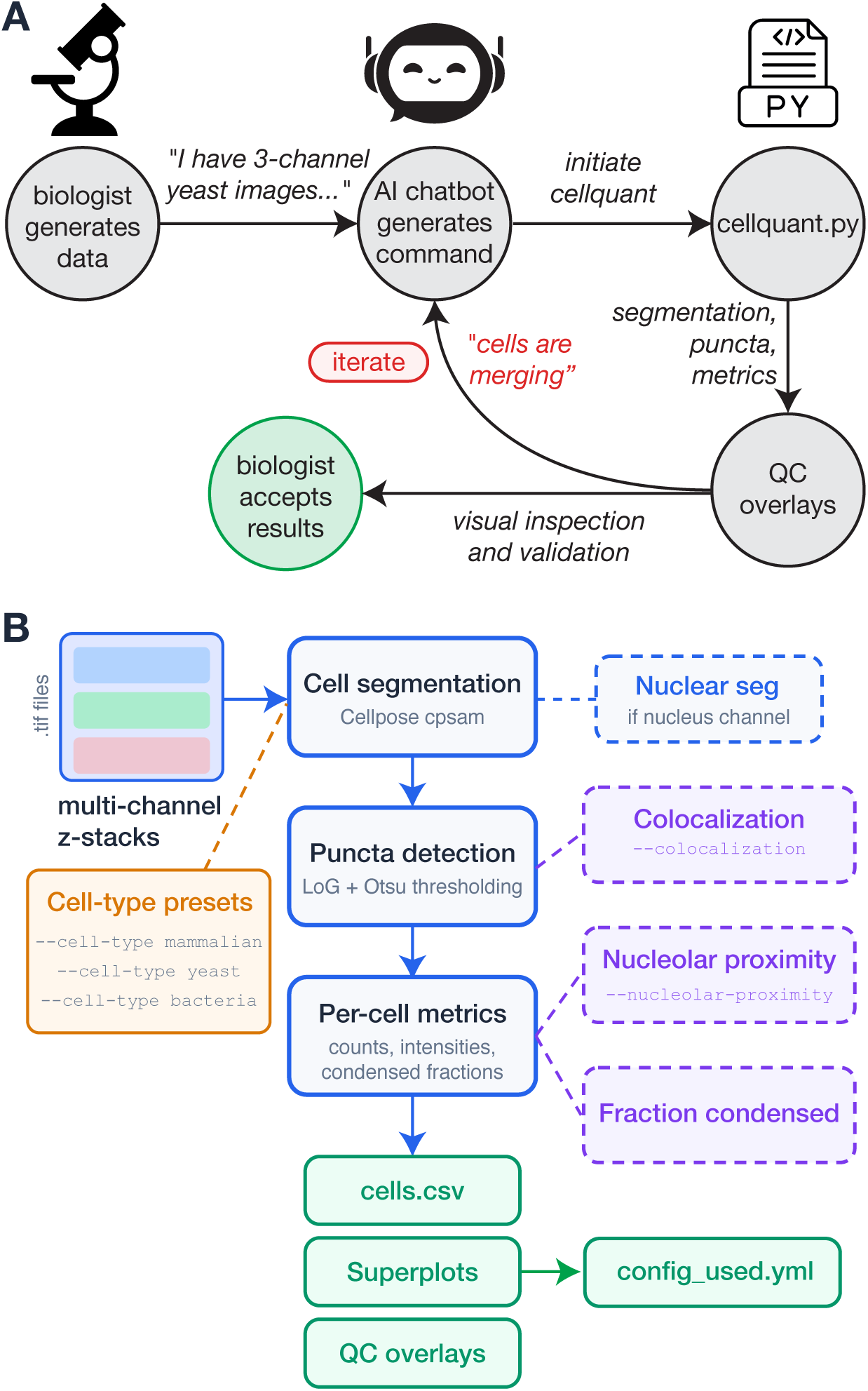
Workflow and cellquant pipeline. **(A)** A biologist describes their analysis to an AI assistant, which generates an appropriate CLI call for `cellquant`.py that the user can paste into the command line. The pipeline produces segmentation, cell metrics, and QC overlays for the biologist to evaluate. If results are not right, the biologist iterates by describing what went wrong to the chatbot until the results pass QC. **(B)** Schematic of the pipeline architecture. Multi-channel z-stacks are processed through cell segmentation (Cellpose cpsam), optional nuclear segmentation when a channel is declared with the nucleus role, puncta detection (Laplacian-of-Gaussian filtering with Otsu thresholding), and per-cell metric computation. Cell-type presets provide organism-specific defaults for every parameter. Modular analyses (colocalization, nucleolar proximity, fraction condensed) are activated by single command-line flags. Every run writes per-cell metrics (cells.csv), superplots, QC overlays, and a complete record of the resolved parameters (config_used.yml).

Here we present `cellquant`, a single Python script that enables high-dimensional quantification of multi-channel fluorescence images. To make it accessible to biologists with all levels of coding experience, even those who may never have opened a terminal window before, `cellquant` is designed to interoperate with a LLM as an assistant. In conversation with a LLM, the user iteratively designs a single call through a command-line interface (CLI) to set all parameters, using cell-type presets and organism-specific defaults where appropriate. In each run, `cellquant` outputs visual segmentation overlays to enable the users to validate or reject the parameter set, describing to the LLM any problems they observe to generate the next CLI call. We pair the pipeline with a comprehensive tutorial to demonstrate the iterative approach. We validate `cellquant` on arsenite-induced stress granules in human HCT116 cells and temperature-dependent nucleolar reorganization in budding yeast. Along with an AI assistant, `cellquant` lets a biologist without programming experience perform accurate and reproducible quantitative image analysis.

## Results

### A single-script pipeline for multi-channel fluorescence image analysis

We implemented a complete image analysis pipeline from segmentation through statistical visualization in a single Python script, cellquant.py (Figure 1B). The pipeline accepts multi-channel fluorescence images in TIFF, OME-TIFF, ND2, CZI, or LIF format, as either two-dimensional (2D) slices or projections or three-dimensional (3D) z-stacks; dimensionality is auto-detected from the input. For cell segmentation and optional organelle segmentation, `cellquant` utilizes Cellpose [Pachitariu and Stringer, 2022] or Cellpose-SAM in 3D; for puncta detection, `cellquant` deploys 2D or anisotropic 3D Laplacian-of-Gaussian filtering; for colocalization analysis, it computes coefficients from the full voxel distribution when input is 3D; for spatial proximity and volume measurements, it outputs physical units derived from voxel size metadata; for data visualization and statistical analysis, it defaults to the “superplot” convention of showing a raincloud plot of individual cells overlaid with replicate medians and calculates *p*-values from the population-level replicate statistics [Lord et al., 2020]. To ensure reproducible quantification across runs, `cellquant` generates a complete provenance record including the exact CLI invocation along with technical metadata for each run.

To tailor `cellquant` for each dataset, the user configures `cellquant` exclusively through a CLI call. Along with a LLM, the user will specify the identity and role (i.e., organelle marker for segmentation or protein of interest to quantify) of each channel in the image file (e.g., “1:DAPI:nucleus” “2:G3BP1:quantify”). We include empirically determined presets to segment different cell types (--cell-type mammalian, yeast, or bacteria), and any preset value can be overridden by explicit arguments.

### Development of cellquant through vibecoding

We developed the pipeline iteratively using the AI coding assistant Claude Code (Anthropic). Importantly, we did not write any new analytical tools. Rather, we flexibly linked multiple open-source, gold standard software packages into a single script. We initially built the mammalian cell pipeline by describing the desired analysis in natural language and refining based on QC overlay evaluation. To extend to yeast and specify new analysis modalities (nucleolar segmentation, colocalization, spatial proximity), we again described what we wanted in natural language, the AI assembled the tools into the script, and we validated through the same visual QC loop.

Several episodes during development illustrate the iterative nature of the approach. When 3*×* downsampling before Cellpose segmentation, initially implemented for computational efficiency, was applied to yeast images, the smaller cell size meant that downsampling destroyed the spatial information needed for accurate segmentation. However, for mammalian cells the same 3*×* downsampling helped segmentation by suppressing signal variation. To support 3D analysis, we described to the LLM that we wanted to incorporate auto-detection of 2D versus 3D input and have `cellquant` perform 3D segmentation when appropriate, identify puncta via anisotropic Laplacian-of-Gaussian detection, calculate colocalization from the full voxel distribution, and use marching cubes for 3D morphometry. The AI assistant implemented the changes by adding to the existing pipeline without disturbing the validated 2D code, and we iteratively validated segmentation using the visual overlays for quality control.

### Installation, tutorial, documentation

Designed to make the entire workflow accessible without programming experience, the `cellquant` pipeline is distributed as a single Python script alongside an environment specification (environment.yml), documentation, and a tutorial. The tutorial guides users of all computational skill levels through installation and operation, from opening a terminal window through a complete analysis of example datasets with outputs at each step. The tutorial demonstrates how to describe an analysis problem to an AI assistant in enough detail to get a working `cellquant` command and iteratively adjust parameters. We also include a complete CLI reference (CLI_REFERENCE.md), which documents every argument `cellquant` takes with its default value, range, and cell type presets. This reference is structured to be readable by both humans and LLMs, so that a user can paste it into a context window and ask an AI assistant for help.

### Stress granule quantification in human cells

To validate the pipeline on a well-characterized biological system, we analyzed three-channel fluorescence images of HCT116 cells expressing the stress granule markers G3BP1-RFP, G3BP2-RFP, and PABPC1-GFP [Smith and Bartel, 2026], stained with DAPI, and treated with or without 500 µM sodium arsenite (Fig. 2A). Arsenite triggers eIF2*α* phosphorylation, leading to translational arrest and condensation of stalled mRNPs into G3BP1/2-positive stress granules that recruit additional RNA-binding proteins including PABPC1 [Kedersha et al., 2016, Protter and Parker, 2016, Glauninger et al., 2025]. Stress granule formation is driven by G3BP1 and G3BP2, which condense in response to increased cytoplasmic mRNA concentrations [Yang et al., 2020, Guillén-Boixet et al., 2020].

**Figure 2:**
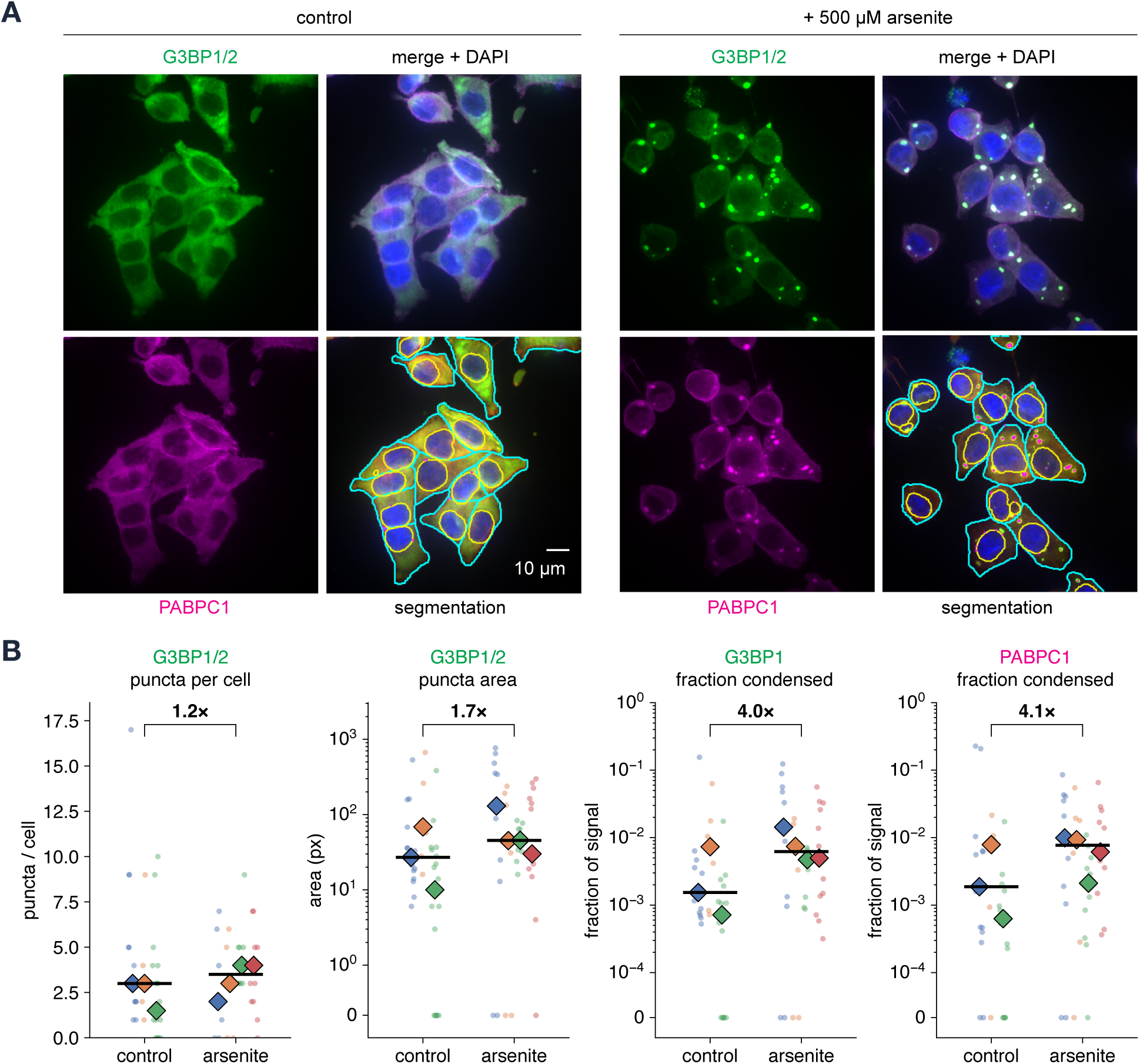
Application of cellquant to stress granules in HCT116 cells. **(A)** Representative maximum intensity projections of HCT116 cells, untreated (left) or treated with 500 µM sodium arsenite for 1 h (right). Top row: G3BP1/2 (green) and merge with PABPC1 (magenta) and DAPI (blue). Bottom row: PABPC1 alone, and the corresponding `cellquant` segmentation showing cell boundaries (cyan), nuclear boundaries (yellow), and detected puncta (magenta). Scale bar, 10 µm. **(B)** Per-cell quantification of G3BP1/2 puncta per cell, G3BP1/2 puncta area, and the fraction of G3BP1 and of PABPC1 signal residing in puncta (*n* = 3 control and 4 arsenite biological replicates, 79 cells). Small points are individual cells colored by biological replicate; large diamonds are replicate medians; the horizontal bar is the median of the replicate medians. Brackets give the fold change (arsenite / control) of the median of replicate medians; *P*-values and Cliff’s *δ* for every metric are in Table S1. The three right-hand panels use a symmetric-log *y* axis so that the *∼*100-fold spread of per-cell values is legible and the cells containing no puncta remain on the plot at zero; the leftmost panel is linear and clipped to the 0.5th–99.5th percentile of the per-cell distribution. Metrics not shown here are given in Fig. S1.

We acquired images as multi-channel z-stacks, with 3–4 biological replicates per condition, and processed them with `cellquant`. Segmentation produced clean cell and nuclear boundaries, and cells treated with arsenite showed visible stress granules containing both G3BP1/2 and PABPC1, while untreated cells showed diffuse cytoplasmic signal (Fig. 2A). Compared to untreated cells, analysis with `cellquant` revealed a modest increase in stress granule size and a substantial increase in the fraction of signal in condensates in both channels in arsenite treated cells (Fig. 2B; Table S1). However, with *n* = 3 versus *n* = 4 replicates, none of the metrics is able to reach conventional significance (the smallest attainable two-sided *p* is 0.057), so we note effect sizes in the superplots.

### Multi-parameter analysis of temperature dependent nucleolar reorganization in yeast

To demonstrate the ability of `cellquant` to generalize across organisms and analysis types, we analyzed three-channel fluorescence images of budding yeast growing at five temperatures from 25°C to 40°C, with 5–8 biological replicates per condition. Cells co-expressed Tif6-Halo, a 60S ribosome biogenesis factor labeled with JF646, Nsr1-mScarlet-I, the yeast nucleolin homolog and nucleolar marker, and Sis1-mVenus, a J-domain protein Hsp40 co-chaperone that forms condensates during heat shock and associates with the nucleolar periphery [Ali et al., 2023, Feder et al., 2021]. Prior to imaging, we cultured cells for six hours at each temperature to allow them to physiologically adapt to the extent possible (Fig. 3A).

**Figure 3:**
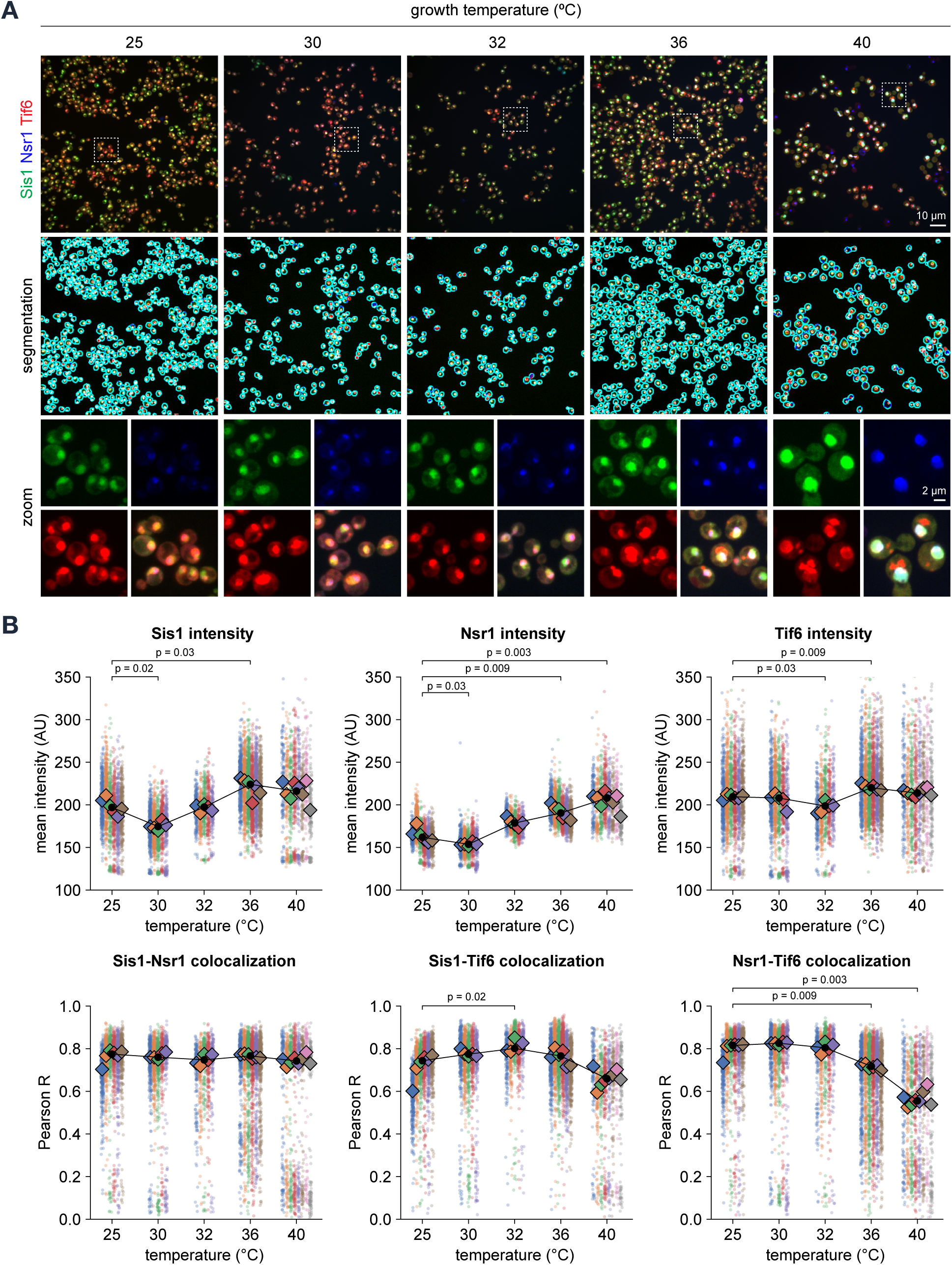
Application of cellquant to temperature dependent nucleolar reorganization in budding yeast. (A) Representative fields at each growth temperature. Top: merge of Sis1-mVenus (green), Nsr1-mScarlet-I (blue), and Tif6-HaloTag/JF646 (red). Middle: `cellquant` cell segmentation. Bottom: magnified views of the boxed regions showing each channel separately and merged. Scale bars, 10 µm (field) and 2 µm (zoom). **(B)** Per-cell quantification across 8,309 cells from 30 biological replicates (6/5/5/6/8 at 25/30/32/36/40°C). Top row: mean cell intensity for each marker. Bottom row: Pearson *R* colocalization for all three channel pairs. Small points are individual cells colored by replicate; large diamonds are replicate medians; the black line joins the median of replicate medians. Brackets show Bonferroni-corrected *P*-values against 25°C where *<* 0.05. Within each family the *y* axes are shared—100–350 AU for intensity, 0–1 for Pearson *R*—so panels can be read against one another; a small number of cells falls outside these ranges and is clipped, while every replicate median lies inside them. Metrics not shown here are given in Fig. S2.

To segment yeast cells, the pipeline linked 2D Cellpose masks across the z-stack using a composite sum of all three channels, since no dedicated membrane or nuclear stain was available.

We filtered out debris and merged cell clusters using volume constraints. The pipeline generated nucleolar masks via Otsu thresholding of the Nsr1 signal.

The pipeline captured a complex set of temperature-dependent changes (Fig. 3B; Table S2; Table S3; Fig. S2). Cell volume increased steadily with temperature, from a median of 37 µm^3^ at 25°C to 65 µm^3^ at 40°C (*p*_adj_ = 0.003), yet nucleolar volume was uncorrelated with temperature. Nsr1 intensity rose progressively across the series (*p*_adj_ = 0.003 at 40°C), while Tif6 intensity was uncorrelated with temperature. Sis1 intensity increased at 36°C and 40°C (*p*_adj_ = 0.017, 0.035). Pairwise colocalization (Pearson’s *R*) computed in 3D revealed that Nsr1-Tif6 colocalization, high at permissive temperatures where both proteins are nucleolar, declined significantly at 36°C and 40°C (*p*_adj_ = 0.009 and 0.003), consistent with Tif6 redistribution upon RiBi repression [Boulon et al., 2010].

### Dimensionality reduction of single cell data resolves subpopulations

The single cell output file (cells.csv) that `cellquant` generated contains 45 non-redundant features per cell, including compartment volumes, mean channel intensities, condensate indices, puncta counts and volumes, fragmentation indices, colocalization coefficients, and nucleolar proximity measurements. To ask how these features collectively change as a function of temperature, we performed principal component analysis (PCA) on the full feature matrix across all five temperature conditions (Fig. 4A).

**Figure 4:**
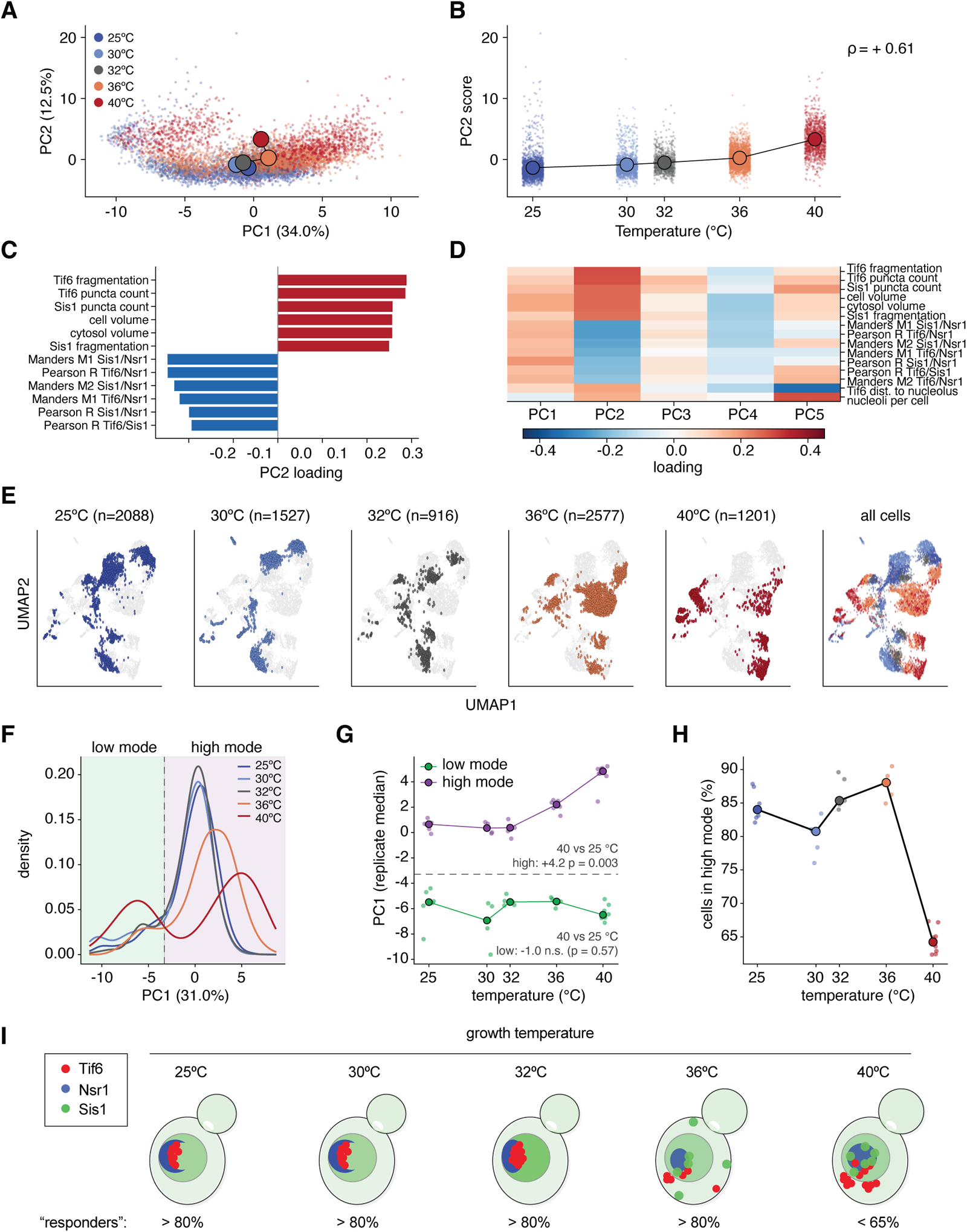
Dimensionality reduction of multi-parameter single cell data reveals nonresponsive subpopulations. **(A)** Principal component analysis over 45 `cellquant` features from 8,309 cells. PC1 versus PC2, cells colored by temperature; large circles are centroids connected in temperature order. **(B)** PC2 score against temperature. **(C)** The twelve features with the largest absolute PC2 loadings. **(D)** Loadings of the same features across PC1–PC5. **(E)** UMAP embedding of the same feature matrix, shown per temperature against the full population in grey, and merged at right. **(F–H)** The population resolves into two modes along PC1 and heat affects only one of them. **(F)** PC1 density by temperature; the dashed line is the antimode of a two-component Gaussian fit (PC1 = *−*3.29; Ashman’s *D* = 2.62), which defines the low and high modes. The second mode is a shoulder at 25°C and resolves into a separate peak at 30°C and above. **(G)** Position of each mode against temperature, as replicate medians. From 25 to 40°C the high mode travels +4.2 PC1 units (*p*_adj_ = 0.003) while the low mode does not move (*−*1.0, *p*_adj_ = 0.57). **(H)** Percentage of cells in the high mode, stable through 36°C and then falling from 84% to 64% at 40°C (*p*_adj_ = 0.003). Panels F–H apply a 10 µm^3^ minimum cell-volume filter (7,569 of 8,309 cells) so that the mode split cannot be attributed to sub-cellular segmentation fragments, which is why PC1 accounts for 31.0% of the variance in panel F against 34.0% in panel A; panels A–E use all cells. **(I)** Cartoon model. With increasing temperature Tif6 (red) disperses from the nucleolus into cytoplasmic/vacuolar puncta, Sis1 (green) forms puncta and moves away from the nucleolar periphery, and Nsr1 (blue) marks the enlarged nucleolus. Above 36°C an increasing fraction of cells are non-responders.

The top mode of variation in the dataset (PC1) does not correlate with temperature, but the second mode (PC2) does (*ρ* = +0.61). The features loading most strongly on PC2 are Tif6 fragmentation and puncta count, Sis1 puncta count, and cell volume in the positive direction, with the colocalization coefficients in the negative direction (Fig. 4C–D). Uniform manifold approximation and projection embedding (UMAP; McInnes et al. 2018), which includes information from deeper modes of variation, showed each temperature occupying a distinct but contiguous region of cell state space (Fig. 4E).

Following statistical decomposition, the single cell distributions revealed two distinct subpopulations along PC1 (Fig. 4F; two-component Gaussian fit, Ashman’s *D* = 2.62, antimode at PC1 = *−*3.29). The low-PC1 mode is largely unresolved from the distribution at 25°C and separates into a distinct peak at 30°C and higher. The high-PC1 mode moves +4.2 PC1 units from 25°C to 40°C (*p*_adj_ = 0.003) while the low-PC1 mode does not move (*p*_adj_ = 0.57) (Fig. 4G). For this reason, we refer to the high-PC1 mode as the “responsive” and the low-PC1 mode as the “non-responsive” populations. Responsive cells constitute *≥* 80% of cells from 25°C to 36°C, but only 64% of cells respond at 40°C (*p*_adj_ = 0.003; Fig. 4H). At 40°C, the cells that continue to respond do so at a higher magnitude than the responders at 36°C, but the population fraction of responders decreases.

These analyses support the following model for reorganization of the nucleolus as a function of temperature in yeast (Fig. 4I). At permissive temperatures, Tif6 and Nsr1 colocalize within the nucleolus and Sis1 is largely diffuse in the nucleoplasm and cytosol. As temperature rises, Tif6 forms puncta and separates from Nsr1, and Sis1 forms puncta away from the nucleolus. An increasing fraction of cells does not respond along this axis at 40°C. This heterogenous reorganization is apparent because `cellquant` enabled us to measure colocalization, morphometrics, and spatial proximity for 8,309 single cells in a single run.

## Discussion

Here we present `cellquant`, an image analysis pipeline that enables researchers with all levels of coding experience to accurately and reproducibly quantify fluorescence images. By working alongside an AI assistant to develop a specific CLI call relevant for each experiment and iterating by visually assessing the outputs and refining the call, we designed `cellquant` to operate in a “vibecoder” loop to customize each analysis while leaving the vetted and versioned codebase unchanged.

Several other groups have explored LLM-mediated image analysis, and they take different approaches. Omega from the Royer laboratory integrates a LLM into napari, allowing the model to generate analysis code within the napari environment [Royer, 2024], and the Fiji community (fiji-llm, available at https://github.com/fiji/fiji-llm) has added a similar feature to Fiji/ImageJ. These tools rely on the LLM to generate analysis code de novo for each run, which allows the user to ask for any analysis the LLM can write. By contrast, the analysis code is fixed in `cellquant`, and the role of the LLM is to help the user generate the specific CLI for their analysis.

The yeast temperature series analysis showcases the high-dimensional measurements `cellquant` can produce. In one experiment, we revealed coordinated changes in cell size with Sis1, Tif6, and Nsr1 re-localization across the temperature series that we quantified all with a single CLI call; we only had to generate eleven lines of specific code. In CellProfiler, this would require the user to construct a custom pipeline, requiring either programming expertise or extensive knowledge of how to use the CellProfiler graphical user interface.

By developing `cellquant`, we in no way suggest that AI can replace computational biologists. Rather, we argue that for standard analyses like segmenting cells and puncta and quantifying colocalization and spatial organization, the combination of a single CLI tool and an AI assistant is a powerful and enabling approach for a biologist to perform rigorous, reproducible image analysis. The simple, modular design of `cellquant` will allow us to extend the platform to additional analysis types, including time courses for live imaging and plate-based formats for high-throughput screens. We will also add more presets for additional cell types and organisms as we establish validated parameters.

More broadly, we envision `cellquant` as a template for “standardized vibecoding”; i.e., a trustworthy codebase that curates the best available open source tools and compiles them for easy and reproducible use by experimentalists working with a LLM to help them customize the analysis. This approach generalizes beyond imaging and could be beneficially applied to any multi-step computational pipeline that currently limits the ability of biologists to analyze their own data, or that may limit their imaginations from planning experiments in the first place for fear of processing the data.

## Limitations of the study

The pipeline supports both 2D and 3D analysis, but it does not currently support time-lapse analysis. Puncta detection assumes approximately spherical (3D) or circular (2D) objects and may not perform well on elongated or irregular shapes. Colocalization analysis uses global Costes thresholding, which may not be optimal for heterogeneous cell populations. The pipeline does not currently support batch parallelization, processing images sequentially. For large screening datasets this may be a practical limitation.

The workflow depends on two external factors beyond the user’s control: the quality of the underlying Cellpose segmentation model and the quality of the AI coding assistant. Cellpose performance varies across cell types and imaging conditions, and the results are only as good as the segmentation. Similarly, AI assistants improve (and occasionally regress) with model updates, meaning that the same description may produce different parameter suggestions. The user’s ability to evaluate QC overlays and recognize when results are biologically incorrect remains the most important safeguard.

## Materials and Methods

### Cell culture and treatment

#### Mammalian cells

HCT116 cells stably expressing G3BP1-RFP, G3BP2-RFP, and PABPC1-GFP [Smith and Bartel, 2026] were cultured in DMEM supplemented with 10% FBS at 37°C with 5% CO_2_. For stress granule induction, cells were treated with 500 µM sodium arsenite for 1 hour or left untreated (vehicle control). Cells were fixed with 4% paraformaldehyde and 4% sucrose, quenched with 125 mM glycine, and stained with DAPI as described [Ali et al., 2023]. G3BP1-RFP, G3BP2-RFP, and PABPC1-GFP signals are from the stably expressed fluorescent fusion proteins.

#### Yeast cells

Budding yeast (*Saccharomyces cerevisiae*, W303 background) expressing endogenously tagged Sis1-mVenus, Tif6-HaloTag, and Nsr1-mScarlet-I (strain construction as described in Ali et al. 2023) were grown to mid-log phase at 30°C in synthetic complete medium. Tif6-HaloTag was labeled with Janelia Fluor 646 HaloTag ligand (JF646, 1 µM) as described [Ali et al., 2023]. For the temperature series, aliquots were shifted to 25, 30, 32, 36, or 40°C for 6 hours starting from a pre-grown log-phase culture at 30°C. Cells were fixed in 1% paraformaldehyde as described [Garde et al., 2024].

#### Image acquisition

All images were acquired on a Nikon SoRa spinning disk confocal microscope (63*×* objective) at the University of Chicago Integrated Light Microscopy Core (RRID: SCR_019197). Exposure time was 100 ms per channel for all acquisitions. Images were collected as z-stacks. For HCT116 cells, z-stacks consisted of 10 slices at 0.50 µm step size; the lateral pixel size was 0.094382 µm/pixel (1192 *×* 1200 pixels = 112.5 *×* 113.3 µm field of view). For yeast, z-stacks consisted of 71 slices at 0.23 µm step size; the lateral pixel size was 0.10571 µm/pixel (1192 *×* 1200 pixels = 126.0 *×* 126.9 µm field of view). The two experiments used different magnification-changer settings, which accounts for the different lateral pixel sizes.

#### Image analysis pipeline

All image analysis was performed using cellquant.py (version 1.1.0, available at https://github.com/davidpincus/cellquant). Two-dimensional analyses were run locally on a MacBook Pro (Apple M-series processor) using CPU-mode Cellpose, owing to MPS GPU incompatibility with the cpsam Transformer model. The three-dimensional yeast analysis was run on the University of Chicago Midway3 compute cluster, also in CPU mode.

### Mammalian cell analysis

Images were processed with the following command:

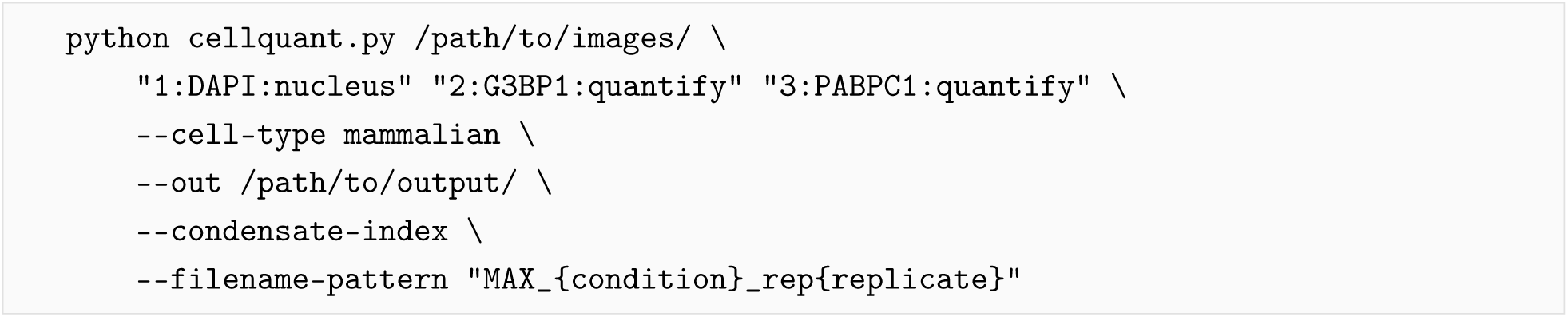

Key parameters (from the mammalian preset): Cellpose cpsam model, 3*×* downsampling, automatic cell diameter estimation, puncta detection in the cytoplasmic compartment (cell minus nucleus).

### Yeast cell analysis

Images were processed with the following command:

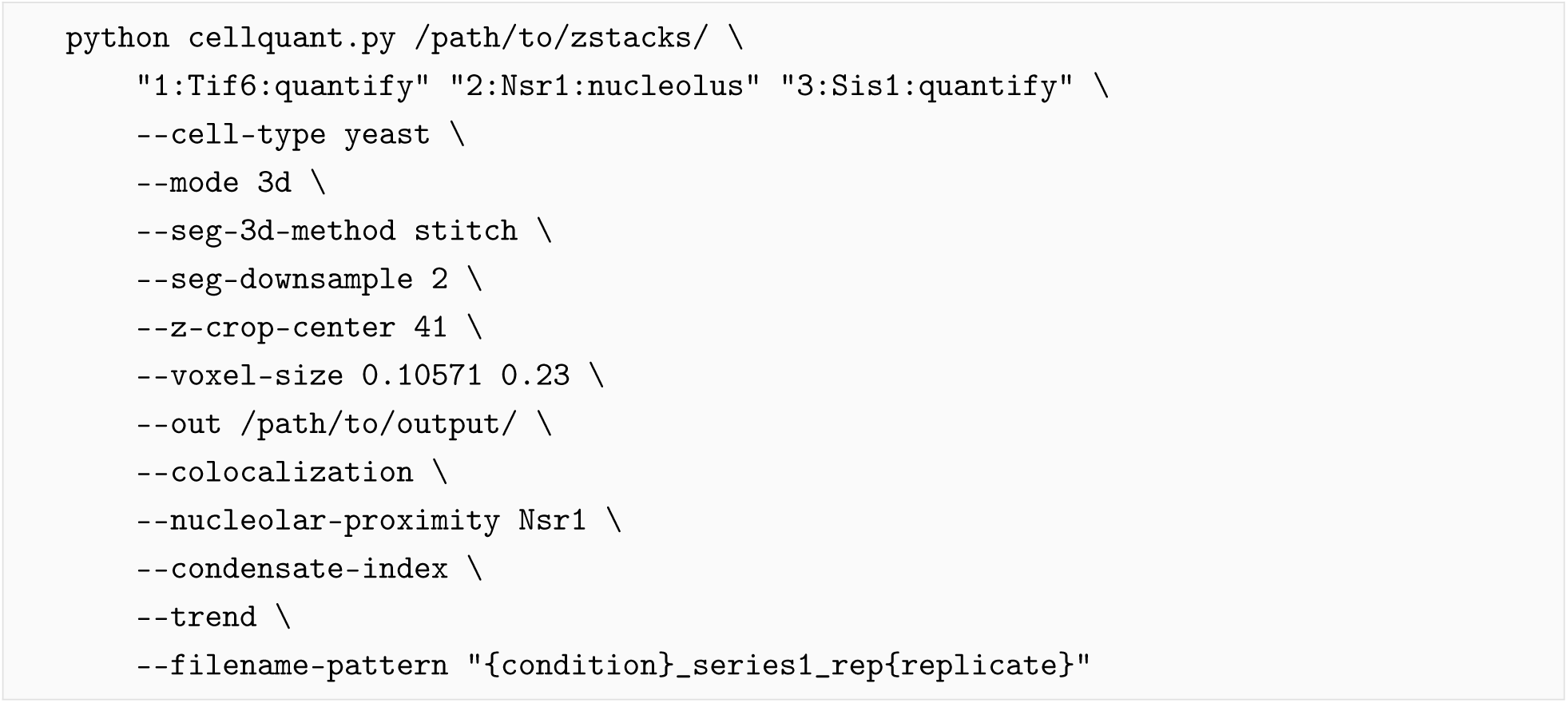

The input is the unprojected z-stack for each replicate. The lateral and axial voxel sizes passed on the command line are those recorded by the instrument in the OME metadata of these files (0.10571 and 0.23 µm), so the run satisfies the 1% agreement check described below rather than overriding it; passing them explicitly makes the analysis reproducible from the deposited images alone. Colocalization is computed natively on the z-stack, and a run of this command on maximum-intensity projections is refused by `cellquant` (see *Three-dimensional segmentation and colocalization*). Key parameters (from the yeast preset, except where set above): Cellpose cpsam model, 2*×* segmentation downsampling, cell diameter 40 pixels, composite-channel segmentation input, cell area filtering (200–5,000 pixels), whole-cell puncta detection, and a 41-plane axially centred crop of each stack. `cellquant` exposes the principal Cellpose parameters (flow threshold, cell-probability threshold, and cell diameter) as CLI flags, providing an alternative to post-hoc area filtering for users who prefer to tune segmentation directly.

### Three-dimensional segmentation and colocalization

Pearson’s correlation coefficient and Manders’ coefficients (M1, M2) are defined on the joint voxel intensity distribution of two channels and require that distribution to reflect the true spatial structure of the imaged volume. When a z-stack is first reduced to a maximum intensity projection (MIP), signal contributions from non-overlapping z-planes are projected onto the same XY position, biasing both Pearson and Manders values in ways that depend on stack depth and the axial distribution of the markers. For 3D inputs, `cellquant` computes colocalization natively on the full z-stack with Costes automatic thresholding [Costes et al., 2004] applied per channel in three dimensions. Colocalization on projections is disabled by default: it requires the explicit --allow-2d-colocalization flag, and any value so produced is stamped as projection-derived in the provenance record. Three-dimensional segmentation itself is performed in one of two modes (selected by --seg-3d-method): a slice-stitching mode applying 2D Cellpose per Z-plane with intersection-over-union-based linkage (faster, more permissive), or a fully volumetric mode using Cellpose-SAM with anisotropy-aware processing (more conservative, typically better for densely sampled stacks). All three cell-type presets default to the slice-stitching mode, at an intersection-over-union threshold of 0.65 for yeast and 0.4 for mammalian cells and bacteria.

### Fragmentation indices

For diffuse or borderline-condensed signal where discrete puncta detection is brittle, `cellquant` reports two complementary threshold-handling regimes of a per-cell fragmentation index per channel: a simple Otsu-thresholded connected-component count (fragmentation_index_simple) and a threshold-free integral over a swept threshold range (fragmentation_index_persistence). The persistence variant provides a measurement that does not depend on a single threshold choice and is therefore more robust when the same puncta-detection settings are applied across heterogeneous conditions.

### Statistical analysis

One convention is used throughout. Per-cell measurements are summarized at the replicate (image) level using medians; the replicate median is the unit of statistical analysis [Lazic et al., 2018], and reported values are the median of those replicate medians. All comparisons use the two-sided Mann-Whitney *U* test as implemented by scipy.stats.mannwhitneyu, which evaluates the exact null distribution at these sample sizes when the ranks are untied and falls back to the normal approximation otherwise. This is the same call `cellquant` makes internally, so every *p*-value reported here is reproduced by re-running the pipeline. At *n* = 3–8 replicates the exact and approximate forms are not interchangeable, so a single implementation is used throughout rather than mixing the two. For the two-condition HCT116 comparison there is a single test per metric and no correction is applied; note that with *n* = 3 and *n* = 4 replicates the smallest attainable two-sided *p* is 0.057. For the multi-condition yeast temperature series (*n* = 5–8 replicates per condition), each temperature is compared to the 25°C baseline using the –reference-condition flag and *p*-values are Bonferroni-corrected across the four temperature comparisons; only comparisons reaching *p*_adj_ *<* 0.05 are annotated in figures. Superplots display per-cell data as violin plots with replicate medians marked as diamonds and individual cells as colored scatter points.

Yeast colocalization coefficients were computed natively on the z-stacks for 30 replicates (6, 5, 5, 6 and 8 at 25, 30, 32, 36 and 40°C), summarized as replicate medians, and compared against the 25°C baseline by two-sided Wilcoxon rank-sum test with Bonferroni correction across the four temperature comparisons.

For dimensionality reduction, the per-cell feature matrix from cells.csv was standardized (zero mean, unit variance per feature) prior to PCA using sklearn.decomposition.PCA. Features with *>*10% missing values were excluded. UMAP embedding used umap-learn with n_neighbors=15 and min_dist=0.1.

### Software and reproducibility

The complete analysis environment is specified in environment.yml and includes Python 3.11–3.12, Cellpose (*≥*4.0, providing both Cellpose and Cellpose-SAM models), scikit-image (*≥*0.24), numpy (*≥*1.24, *<*2.0), opencv-python-headless (*<*4.10), pandas, matplotlib, scipy, PyYAML, and bioio with format-specific readers for ND2, CZI, and LIF input. `cellquant` also ships with an inline PEP 723 script header so that uv run `cellquant`.py resolves and installs all dependencies on first invocation, eliminating the conda environment step for users with uv installed. All code, example data, and expected outputs are available at https://github.com/davidpincus/cellquant, and the version used here (v1.1.0) is archived at Zenodo (DOI: 10.5281/zenodo.21830047). Full imaging datasets are deposited at Zenodo (DOI: 10.5281/zenodo.21843280).

### Metadata handling and provenance

`cellquant` reads image metadata at run time and propagates it through the pipeline rather than assuming pixel-unit defaults. For OME-TIFF inputs, voxel sizes in X, Y, and Z (in microns) and channel names are parsed from the Pixels and Channel elements of the embedded XML; for ImageJ-style TIFFs, voxel sizes are read from the resolution and spacing tags. When voxel size metadata is missing or values fall outside plausible microscopy ranges (XY voxel *<* 0.01 or *>* 10 µm; Z voxel *<* 0.05 or *>* 50 µm; Z voxel smaller than XY voxel), `cellquant` aborts a three-dimensional run rather than substituting a default, and names the flag that supplies the missing value (--voxel-size, or --assume-isotropic to proceed without one). A voxel size that disagrees with the file metadata by more than 1% is likewise a hard error unless --override-metadata is given; both values and the source of the resolution actually used are recorded in provenance.json. Channel names from OME metadata are cross-checked against the user-supplied --channels specification; mismatches generate a warning so that out-of-order channel declarations are surfaced rather than silently propagated to downstream measurements. Non-TIFF formats (ND2, CZI, LIF) are read through the bioio backend, which exposes the same physical-unit and channel-name metadata through its API. Every `cellquant` run writes a provenance.json sidecar alongside its outputs. The sidecar records the `cellquant` version, the exact CLI, the SHA-256 checksum and modification time of every input file, the OME channel names and voxel sizes read from each input, the Cellpose and Cellpose-SAM model checkpoints used, and pinned versions of every key Python dependency (Python, numpy, scipy, scikit-image, pandas, matplotlib, PyYAML, tifffile, OpenCV, PyTorch, and Cellpose). Together with the config_used.yml configuration record that `cellquant` has always written, the provenance file is sufficient to reconstruct any analysis given the same inputs. The motivation is FAIR alignment of derived measurements [Wilkinson et al., 2016]: a colocalization value or puncta count in cells.csv is only as findable, accessible, interoperable, and reusable as the record of how it was produced. The metadata conventions `cellquant` follows align with current community recommendations for bioimaging publication [Bialy et al., 2024].

## Acknowledgments

We thank Jarrett Smith, David P. Bartel, Joshua Melamed, and D. Allan Drummond for providing the HCT116 cells expressing G3BP1-RFP, G3BP2-RFP, and PABPC1-GFP [Smith and Bartel, 2026]. We thank Madeline Herwig and Jenny Krystosek for beta testing `cellquant` and providing valuable feedback. The `cellquant` pipeline was developed iteratively using the AI coding assistants Claude and Claude Code (Anthropic), which also assisted with repository setup and manuscript preparation. We acknowledge the University of Chicago Integrated Light Microscopy Core (RRID: SCR_019197) for microscopy support. This work was supported by NIH grant RM1 GM153533 and NSF QLCI QuBBE grant OMA-2121044 to D.P.

## Author Contributions

A.N. generated the mammalian stress granule imaging data. A.A. generated the yeast strain and performed the temperature series imaging data along with L.C.W. D.P. conceived the project, developed the `cellquant` pipeline and documentation, performed all analyses, and wrote the manuscript with input from all authors.

## Declaration of Interests

The authors declare no competing interests.

## Data Availability

All code and documentation are available at https://github.com/davidpincus/cellquant. The version of cellquant used here (v1.1.0) is archived at Zenodo (DOI: 10.5281/zenodo.21830047). Full imaging datasets are deposited at Zenodo (DOI: 10.5281/zenodo.21843280).

## Supplemental Materials

**Figure S1:**
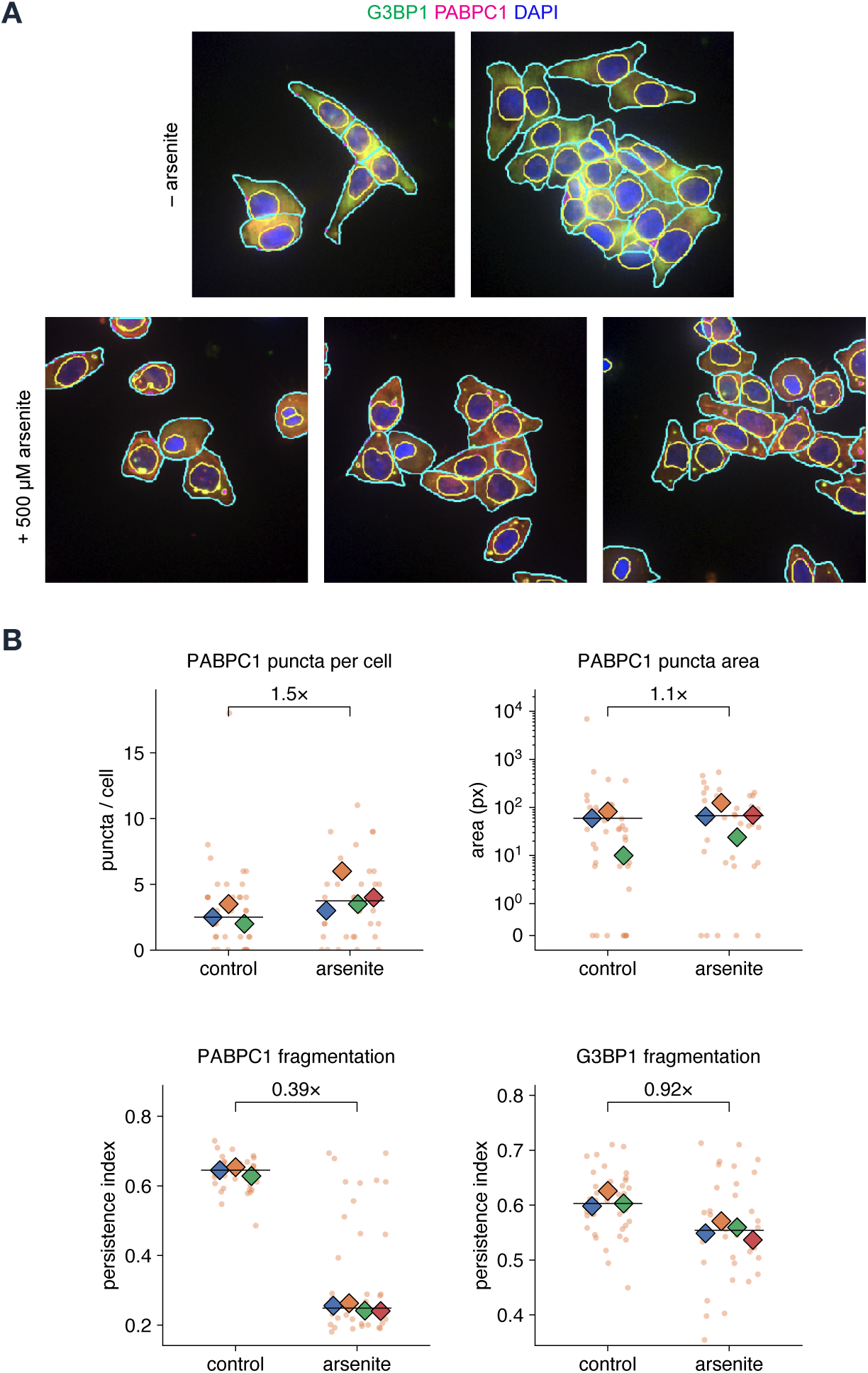
Additional metrics from HCT116 stress granule quantification. **(A)** Maximum-intensity projections with cellquant segmentation overlays: G3BP1 (green), PABPC1 (magenta), and DAPI (blue), untreated (top) or treated with 500 µM sodium arsenite for 1 h (bottom). Cell boundaries are drawn in cyan, nuclear boundaries in yellow, and detected puncta in magenta. **(B)** Per-cell metrics: PABPC1 puncta per cell and puncta area, and the fragmentation index for both channels. Plotting conventions and statistics are as in Fig. 2B, and brackets give the fold change (arsenite / control). The puncta-area panel uses a symmetric-log *y* axis so that cells containing no puncta remain on the plot at zero. The *fragmentation persistence index* is a threshold-free measure on [0, 1]: the signal is swept across all intensity thresholds and the index integrates how persistently it remains broken into separate objects, so it does not depend on choosing any single threshold. Remaining metrics, including the condensate indices, are given in Table S1.

**Figure S2:**
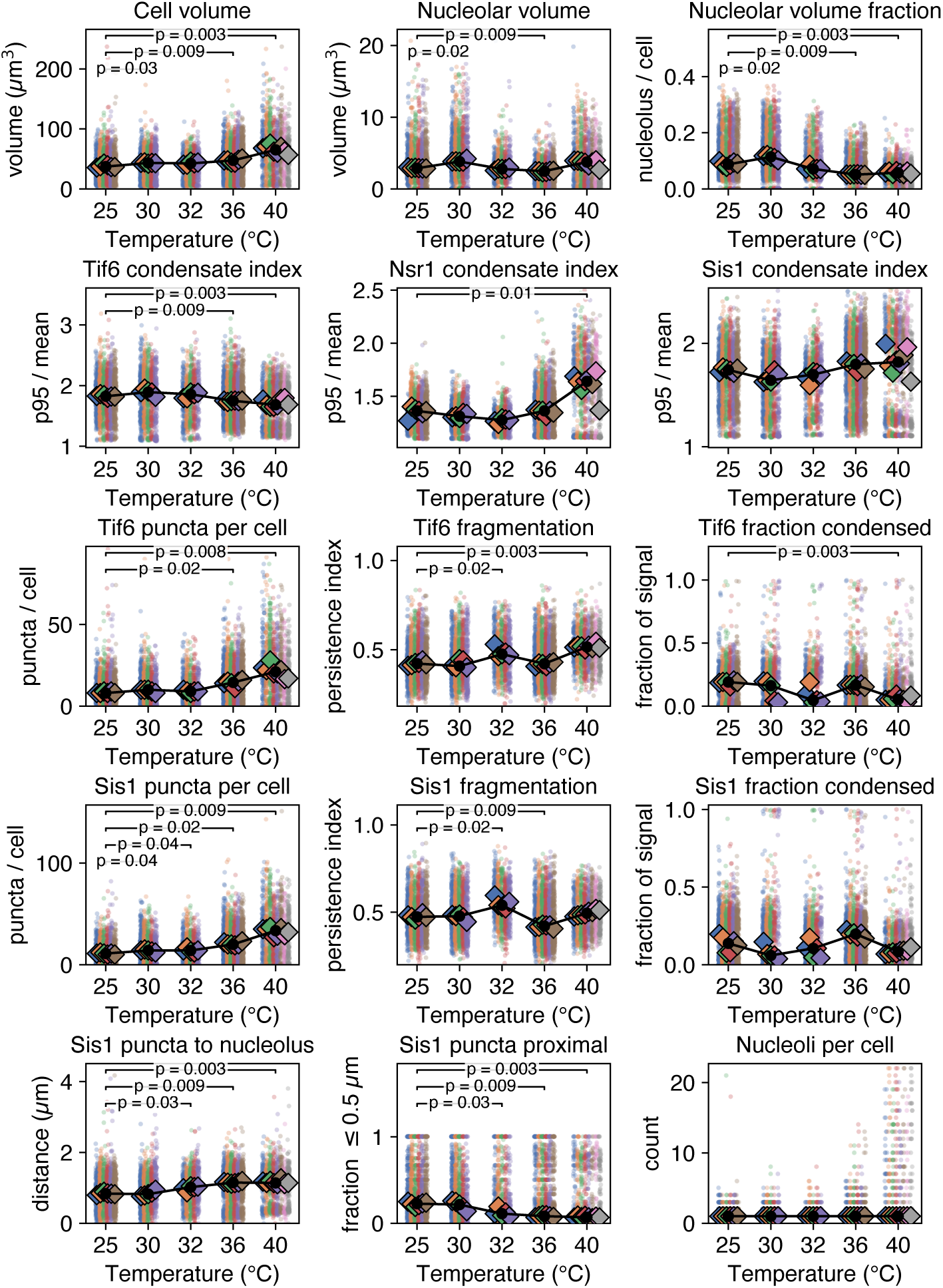
Additional cellquant output for the yeast temperature series. Per-cell metrics cellquant reports for the yeast series that are not shown in Fig. 3B, computed on the z-stacks. All panels use the conventions of Fig. 3B: individual cells as small points colored by replicate, replicate medians as large diamonds, and the black line joining the median of replicate medians across 8,309 cells from 30 biological replicates. Brackets show Bonferroni-corrected *P*-values against 25°C, and only significant comparisons are drawn. Rows, top to bottom: compartment size (cell volume, nucleolar volume, and nucleolar volume as a fraction of cell volume); condensate index for all three markers (the ratio of the 95th-percentile to the mean intensity within each cell, so values *>* 1 indicate non-uniform signal); Tif6 puncta count, fragmentation, and fraction condensed; the same three for Sis1; and nucleolar proximity (mean Sis1 puncta-to-nucleolus distance and the fraction of Sis1 puncta within 0.5 µm of the nucleolus) with nucleoli per cell. Nucleolar area, circularity, eccentricity, and solidity are properties of a two-dimensional projection and have no three-dimensional counterpart; nucleolar volume and volume fraction are reported in their place. Manders’ M1 and M2 for all three channel pairs are omitted because Fig. 3B already reports colocalization for every pair; they are included in the deposited per-cell tables.

**Figure S3:**
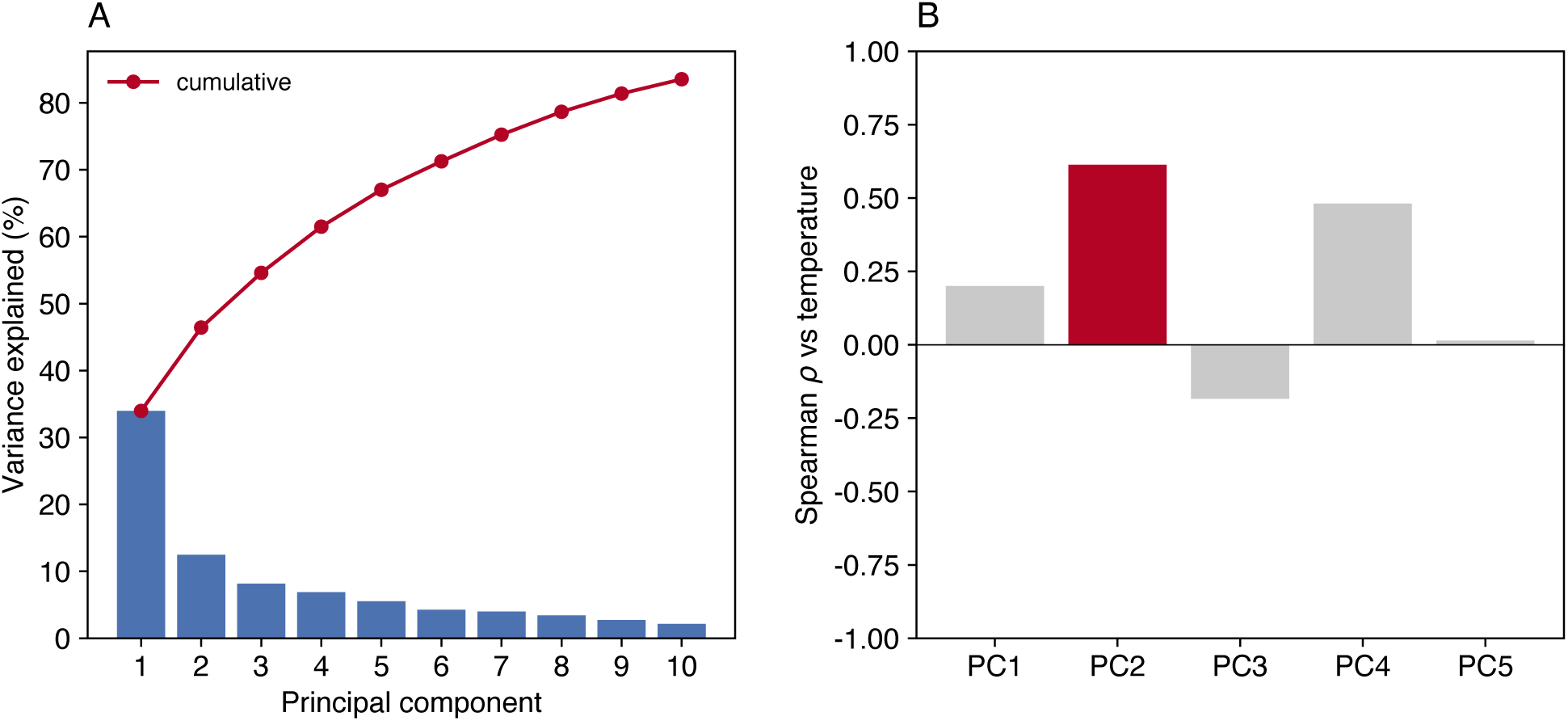
Principal component analysis diagnostics. **(A)** Variance explained by each of the first ten principal components (bars) and the cumulative total (line). PC1 accounts for 34.0% and the first five components for 67.0% of the variance in the 45-feature matrix. **(B)** Spearman correlation between each of the first five components and growth temperature. Temperature is carried by PC2 (*ρ* = +0.61, highlighted), not by PC1 (*ρ* = +0.20), which is why Fig. 4B plots PC2 against temperature.

---

**Table S1:** HCT116 stress granule summary statistics. Values are the median of per-replicate medians. Fold change is arsenite / control; Cliff’s *δ* is the rank effect size belonging to the test performed, bounded [*−*1, 1], where +1 is complete separation. *P*-values are from two-sided Mann-Whitney tests on replicate medians (control: *n* = 3; arsenite: *n* = 4 replicates). At these sample sizes the smallest attainable two-sided *p* is 0.057, so this experiment can establish effect sizes but cannot establish significance; the effect-size columns, not the *P* column, carry the result.

| Metric | Control | Arsenite | Fold change | Cliff's $\delta$ | $P$-value |
| --- | --- | --- | --- | --- | --- |
| G3BP1 puncta per cell | 3 | 3.5 | 1.2× | +0.50 | 0.35 |
| G3BP1 puncta area | 27 | 45.25 | 1.7× | +0.50 | 0.40 |
| G3BP1 fraction condensed | 0.001537 | 0.006206 | 4.0× | +0.67 | 0.23 |
| PABPC1 fraction condensed | 0.001875 | 0.007728 | 4.1× | +0.67 | 0.23 |
| PABPC1 puncta per cell | 2.5 | 3.75 | 1.5× | +0.75 | 0.15 |
| G3BP1 condensate index | 1.421 | 1.466 | 1.0× | +0.50 | 0.40 |
| PABPC1 condensate index | 1.51 | 1.463 | 0.97× | -0.33 | 0.63 |

**Table S2:** Yeast temperature series summary. All metrics computed natively on the z-stacks. Values are the median of per-replicate medians across 5–8 biological replicates per temperature. Nucleolar shape descriptors are properties of a projection and have no three-dimensional counterpart; nucleolar volume and equivalent diameter are reported in their place. Statistical comparisons are given in Table S3.

| Metric | 25°C | 30°C | 32°C | 36°C | 40°C |
| --- | --- | --- | --- | --- | --- |
| *n* cells | 2088 | 1527 | 916 | 2577 | 1201 |
| *n* replicates | 6 | 5 | 5 | 6 | 8 |
| *Condensate metrics* |  |  |  |  |  |
| Sis1 condensate index | 1.74 | 1.64 | 1.70 | 1.80 | 1.82 |
| Sis1 frac. condensed | 0.137 | 0.060 | 0.105 | 0.197 | 0.084 |
| Sis1 puncta/cell | 11.00 | 14.00 | 14.00 | 20.00 | 33.50 |
| Tif6 condensate index | 1.82 | 1.89 | 1.86 | 1.75 | 1.68 |
| Tif6 frac. condensed | 0.191 | 0.162 | 0.046 | 0.167 | 0.053 |
| Tif6 puncta/cell | 8.00 | 10.00 | 9.00 | 14.50 | 21.00 |
| Nsr1 condensate index | 1.36 | 1.31 | 1.28 | 1.36 | 1.64 |
| *Colocalization* |  |  |  |  |  |
| Pearson R (Tif6 vs Sis1) | 0.74 | 0.77 | 0.80 | 0.77 | 0.66 |
| Pearson R (Tif6 vs Nsr1) | 0.82 | 0.83 | 0.80 | 0.72 | 0.56 |
| Pearson R (Sis1 vs Nsr1) | 0.77 | 0.76 | 0.75 | 0.77 | 0.74 |
| Manders M1 (Tif6 vs Sis1) | 0.69 | 1.00 | 1.00 | 1.00 | 1.00 |
| Manders M1 (Tif6 vs Nsr1) | 0.87 | 0.83 | 0.83 | 0.74 | 0.59 |
| Manders M1 (Sis1 vs Nsr1) | 0.84 | 0.82 | 0.81 | 0.76 | 0.68 |
| *Nucleolar proximity* |  |  |  |  |  |
| Sis1 fraction proximal | 0.23 | 0.22 | 0.11 | 0.08 | 0.07 |
| Sis1 mean distance (μm) | 0.84 | 0.82 | 1.02 | 1.15 | 1.14 |
| Tif6 fraction proximal | 0.27 | 0.22 | 0.12 | 0.14 | 0.12 |
| Tif6 mean distance (μm) | 0.81 | 0.90 | 1.01 | 1.10 | 1.07 |
| *Cell and nucleolar size* |  |  |  |  |  |
| Cell volume (μm<sup>3</sup>) | 37.2 | 43.4 | 42.5 | 47.5 | 65.1 |
| Nucleolar volume (μm<sup>3</sup>) | 2.88 | 3.82 | 2.78 | 2.48 | 3.75 |
| Nucleolar eq. diameter (μm) | 1.79 | 2.00 | 1.76 | 1.68 | 1.94 |

**Table S3:**
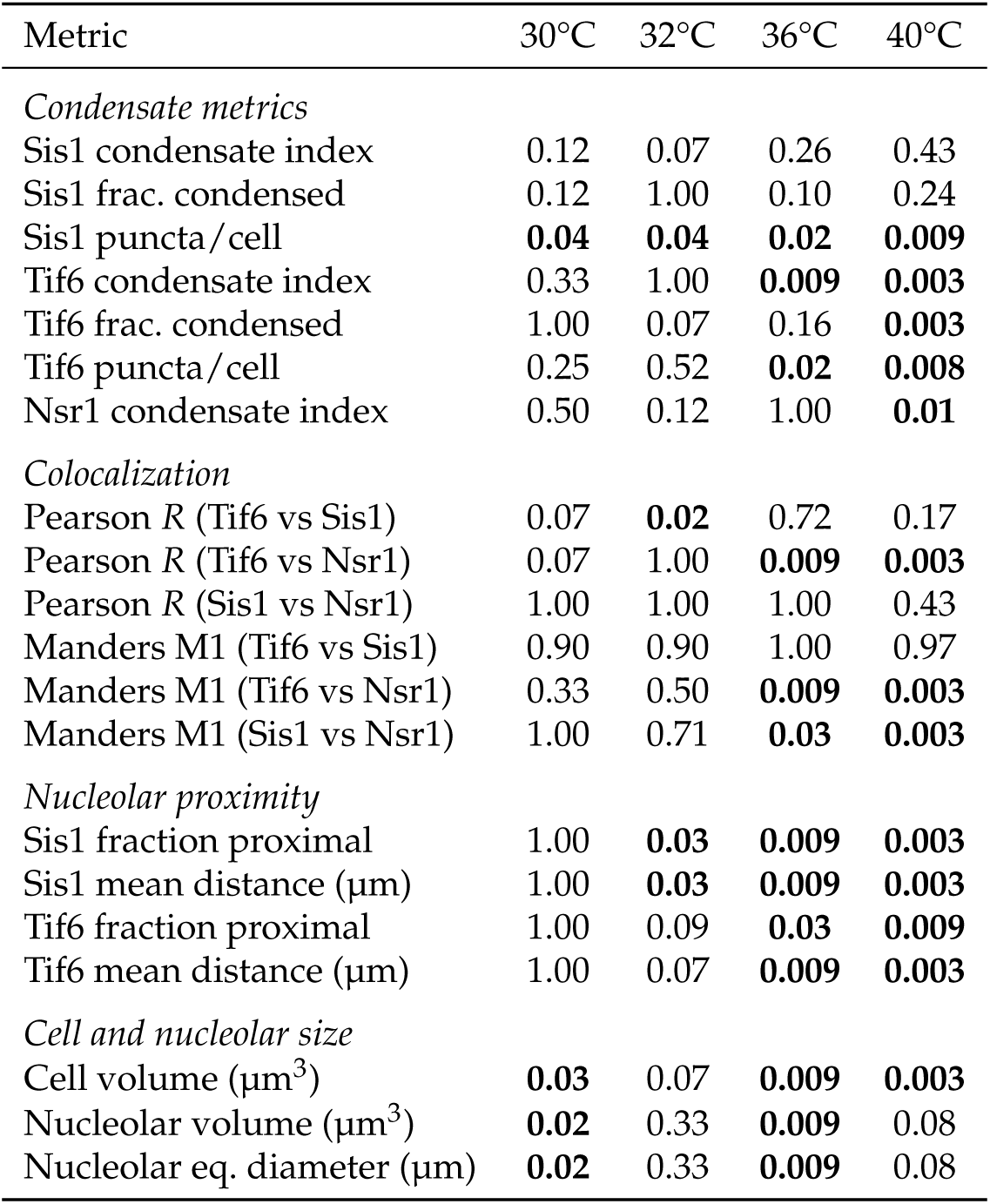
Statistical comparisons for the yeast temperature series. Bonferroni-corrected *p*-values from two-sided Mann-Whitney tests on replicate medians, comparing each temperature to the 25°C reference across the four temperature comparisons. Bold values indicate *p*_adj_ *<* 0.05. The significant 32°C Tif6–Sis1 Pearson comparison is an *increase* over 25°C; every other significant colocalization change is a decrease.

**Table S4:** cellquant parameters. Complete parameter settings used for the HCT116 mammalian and yeast temperature series analyses. Parameters not listed used default values. The HCT116 analysis is two-dimensional and its size parameters are therefore in pixels; the yeast analysis is three-dimensional and its size parameters are in voxels, which are the quantities the three-dimensional code path uses. Because a Z crop was applied to the yeast stacks, cells whose masks touch the top or bottom plane of the cropped volume were excluded so that axially truncated cells did not enter the per-cell measurements.

| Parameter | HCT116 analysis | Yeast analysis |
| --- | --- | --- |
| Analysis mode | 2D (MIP) | 3D (z-stack) |
| Cell type preset | mammalian | yeast |
| Cellpose model | cpsam | cpsam |
| 3D seg. method | n/a | slice stitching, IoU 0.65 |
| Voxel size ($\mu\text{m}$) | n/a (pixel units) | 0.10571 lateral, 0.23 axial |
| Z crop (planes) | n/a | 41, axially centred |
| Seg. downsample | 3× | 2× |
| Cell diameter (px) | auto | 40 |
| Seg. channel | PABPC1 | composite |
| Cell size filter | default | 1,500–200,000 vox |
| Cellprob threshold | 0.0 | –1.0 |
| Puncta channels | G3BP1, PABPC1 | Sis1, Tif6 |
| Puncta compartment | cytosol | whole-cell |
| LoG sigma | 2.0 | 1.5 |
| Puncta min size | 6 px | 4 vox |
| Puncta max size | 10,000 px | 3,000 vox |
| Threshold method | Otsu | Otsu |
| Nuclear req. | 1–4 nuclei | none |
| Nucleus dilation (px) | 3 | 0 |
| Colocalization | no | yes (whole-cell, native 3D) |
| Nucleolar proximity | no | yes (Nsr1, 0.5 $\mu\text{m}$) |
| Plot style | superplot | trend |

## Notes

https://github.com/davidpincus/cellquant

### Competing Interest Statement

The authors have declared no competing interest.

### Summary of Updates

The new version has added 3D image segmentation, co-localization, and distance quantification.

